# Prospectively predicting BPaMZ Phase IIb outcomes using a translational preclinical mouse to human platform

**DOI:** 10.1101/2023.02.16.528876

**Authors:** Qianwen Wang, Janice JN Goh, Nan Zhang, Eric Nuermberger, Rada Savic

**Affiliations:** Department of Bioengineering and Therapeutic Sciences, University of California, San Francisco, San Francisco, California, United States of America; Center for Tuberculosis Research, Department of Medicine, Johns Hopkins University School of Medicine, Baltimore, Maryland, United States of America

## Abstract

Despite known treatments, tuberculosis (TB) remains the world’s top infectious killer, highlighting the pressing need for new drug regimens. To prioritize the most efficacious drugs for clinical testing, we previously developed a PK-PD translational platform with bacterial dynamics that reliably predicted short-term monotherapy outcomes in Phase IIa trials from preclinical mouse studies. In this study, we extended our platform to include PK-PD models that account for drug-drug interactions in combination regimens and bacterial regrowth in our bacterial dynamics model to predict cure at end of treatment and relapse 6 months post-treatment. The Phase III trial STAND, testing new regimen pretomanid (Pa), moxifloxacin (M), and pyrazinamide (Z) (PaMZ), predicted to shorten treatment duration by 2 months was put on hold after a separate ongoing trial showed adding bedaquiline (B) to the PaMZ regimen (SimpliciTB) suggested superior efficacy. To forecast if the addition of B would indeed benefit the PaMZ regimen, we applied an extended translational platform to both regimens. We predicted currently available short- and long-term clinical data well for drug combinations related to BPaMZ. We predict the addition of B to PaMZ will shorten treatment duration by 2 months and be non-inferior compared to control HRZE, both at the end of treatment for treatment efficacy and 6 months after treatment has ended in relapse prevention. Using BPaMZ as a case study, we have demonstrated our translational platform can predict Phase II and III outcomes prior to actual trials, allowing us to better prioritize the regimens most likely to succeed.

## Introduction

Tuberculosis (TB) is a communicable disease caused by the bacillus *mycobacterium tuberculosis* (Mtb). According to the World Health Organization (WHO), TB is a major cause of ill health and leading cause of death worldwide ^1^. While therapies are available for TB treatment, the treatment is lengthy and burdensome thus making it hard to eradicate ^2,3^. Therefore, a huge emphasis has been placed on the need for more effective and shorter regimens. ^4,5^.

The development of novel drug combinations heavily relies on evidence from preclinical efficacy models. Among these, murine TB models are the most commonly employed ^6,7^. Due to the highly evasive and resistant nature of Mtb, TB treatment is often implemented as a cocktail of drugs rather than a single drug to decrease resistance and improve overall efficacy while dosing within safe ranges. New drugs are hence often evaluated in combination with a preexisting backbone of known drugs. Recently, novel nitroimidazole, pretomanid (Pa), was tested with known TB drugs moxifloxacin (M) and pyrazinamide (Z) as a multidrug combination (PaMZ) in Phase III clinical trial STAND (NC-006). However, STAND was halted early and analysis of its results showed PaMZ regimens failed to achieve non-inferiority to HRZE despite efficacy in preclinical combinations evaluated in mice^8^. This highlights the complexity of translating our preclinical efficacy to clinical outcomes.

With diarylquinoline, bedaquiline (B), showing good efficacy in other regimens such as BPaL^9^, the BPaMZ regimen was studied to see if the addition of B would improve its efficacy compared to standard of care. Our lab has previously developed a translational pharmacokinetic-pharmacodynamic (PK-PD) platform that can predict Phase IIa outcomes based on preclinical mouse efficacy. The model incorporates bacterial dynamics with mouse exposure-response relationships and human PK profile to predict Phase IIa outcomes for drugs tested as monotherapy. It was validated with good predictions for nine 1^st^ and 2^nd^ line TB drugs. We would thus like to utilize this powerful platform to further ask if we can predict Phase IIb and III outcomes for drug efficacy in combination using BPaMZ as a case study.

In comparison to monotherapy, understanding the contribution of each individual drug to the overall efficacy makes combination therapy a more complex problem to tackle. In preclinical settings, in vitro checkerboard analysis is commonly used to systematically evaluate drug efficacy of combinations ^10,11^. Carrying out such a large range of doses in animals however is impractical due to the scale and number of combinations required. In animal studies, the new drug (B) is often tested as a dose range with a fixed dose combination (e.g. PaMZ) rather than testing all possible dose ranges for a 4-drug combination. Using a PK-PD model informed way to measure if the addition of B to PaMZ would thus be a more effective, cost-saving method to predict efficacy prior to clinical trial.

Unlike Phase IIa outcomes which measured early bactericidal activity in monotherapy for up to 2 weeks, Phase IIb trials focus on the sterilizing potential of the regimen, using time to culture conversion (TTP), while the proportion of patients without relapse serves as the main outcome of interest in Phase III trials. We thus added a few additional components to extend our translation platform: 1) inclusion of pharmacodynamic drug-drug interactions such as synergism and/or antagonism; 2) drug efficacy in both fast- and slow-replicating bacteria; 3) bacteria regrowth kinetics post-treatment (Figure 1). These additions allow us to predict both short- and long-term efficacy outcomes, which are a better reflection of Phase IIb/III outcomes and real-life treatment outcomes. In the long run, we hope to be able to use this platform to prioritize the most effective drug combination for Phase IIb/III trials.

**Figure 1.**
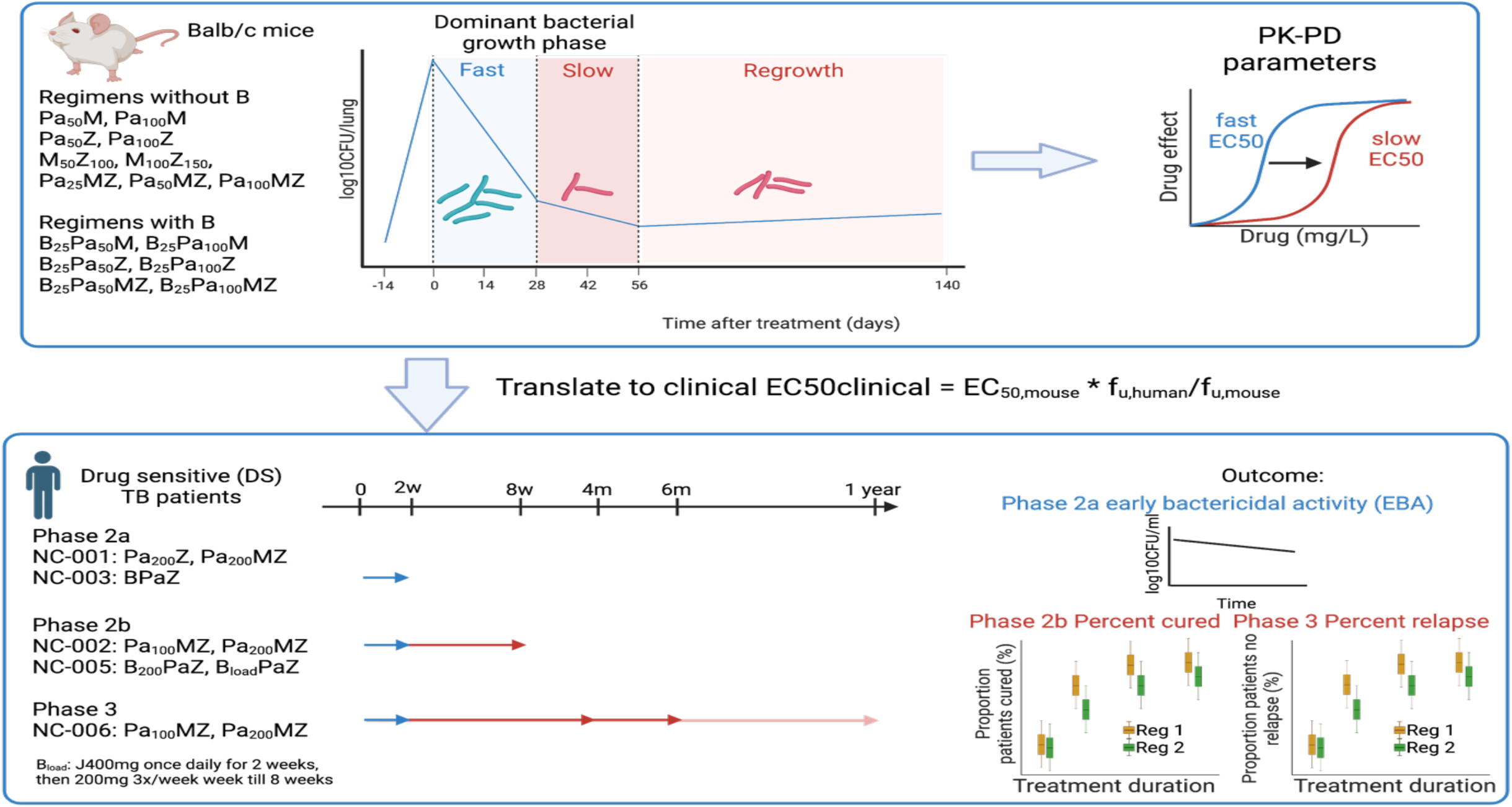
Schematic overview of proposed translational platform for drug combinations. Translational platform for short- and long-term treatment: upper left panel summarized all combinations and regimens that are available in mice. PK-PD mouse models will be built upon drug potency in both fast and slow growth phases and regrowth estimated 3 months post-treatment. Lower left panel describes trials that were finished with reported results, which served as validation of our platform for both long and short-term outcomes.

## Results

### Large database of (B)PaMZ combinations in mice and human

Rich longitudinal preclinical and clinical data were pooled from in-house databases (Figure 2, Figure 3, Table 1) which include: 1) mice longitudinal plasma concentrations (PK data) and CFU counts (PD data) after monotherapy treatments of Pa, M, Z, and B at multiple dose levels; 2) mice longitudinal CFU counts after combinations of PaM, PaZ, PaMZ, BPaM, BPaZ, BPaMZ as well as relapse data 3-month post-treatment; 3) human sputum data in trials including NC-001, NC-002, NC-003, NC-005 and patients time to culture conversion as well as relapse data in trial NC-006. Detailed descriptions of the dataset are in Supplemental Results.

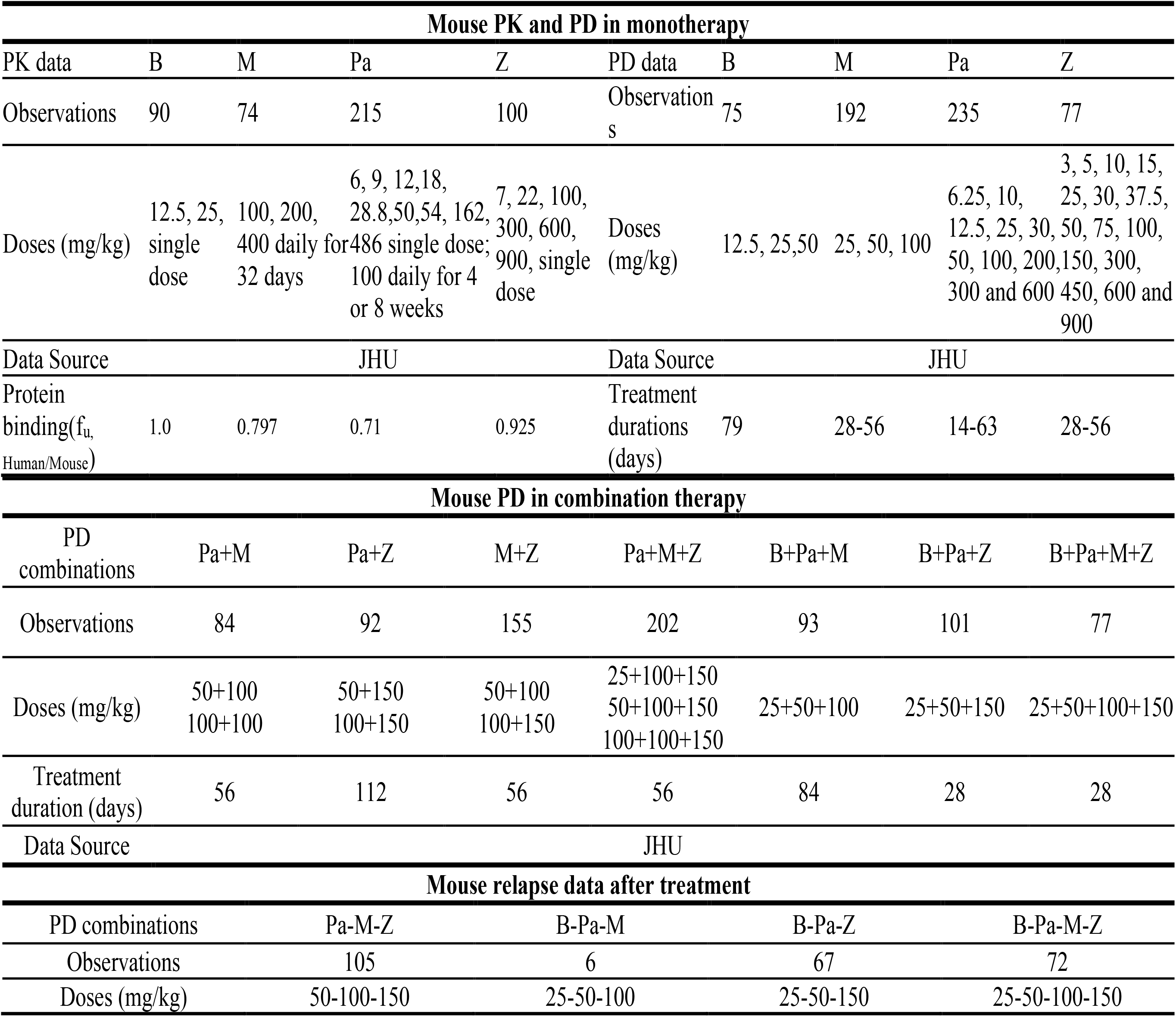
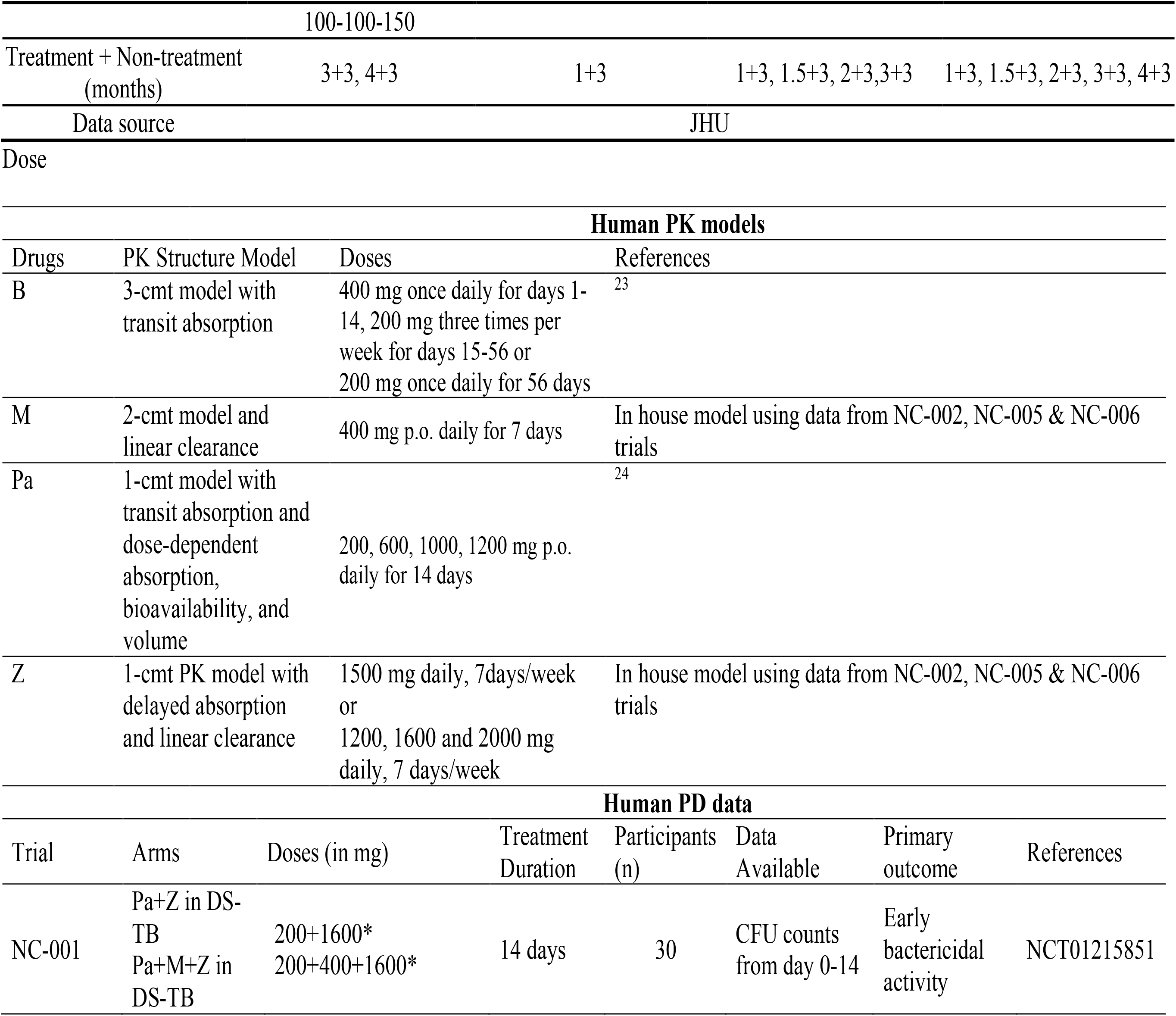
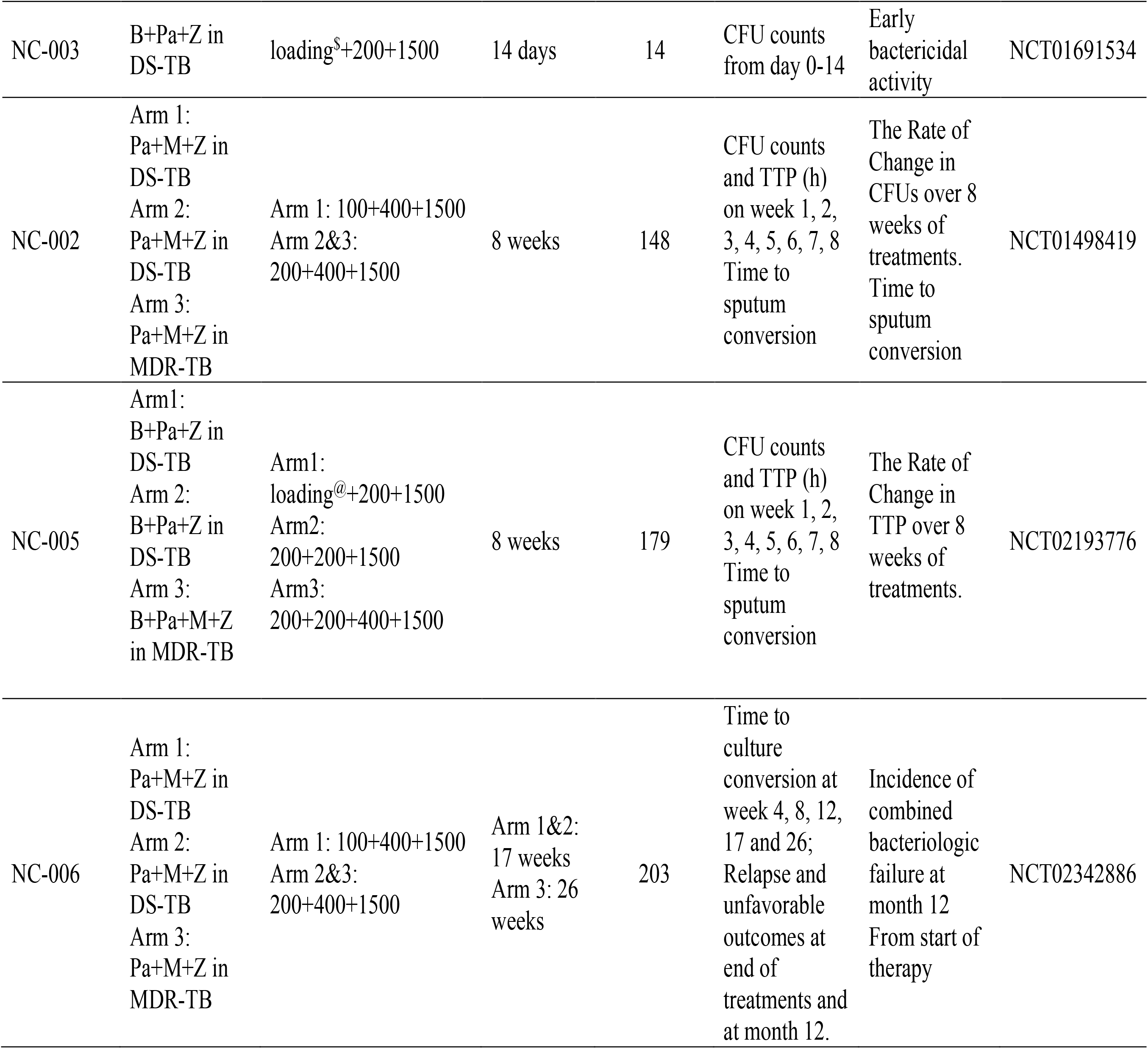
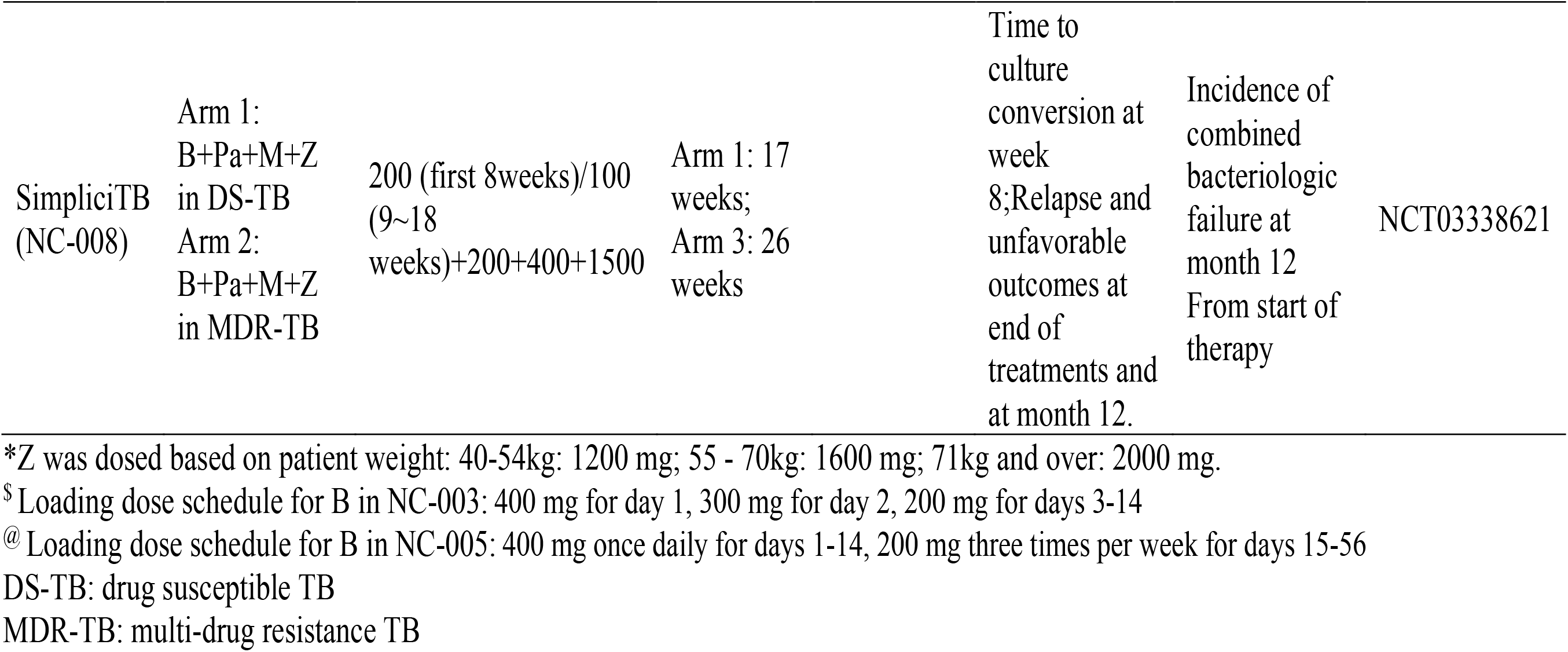

**Figure 2.**
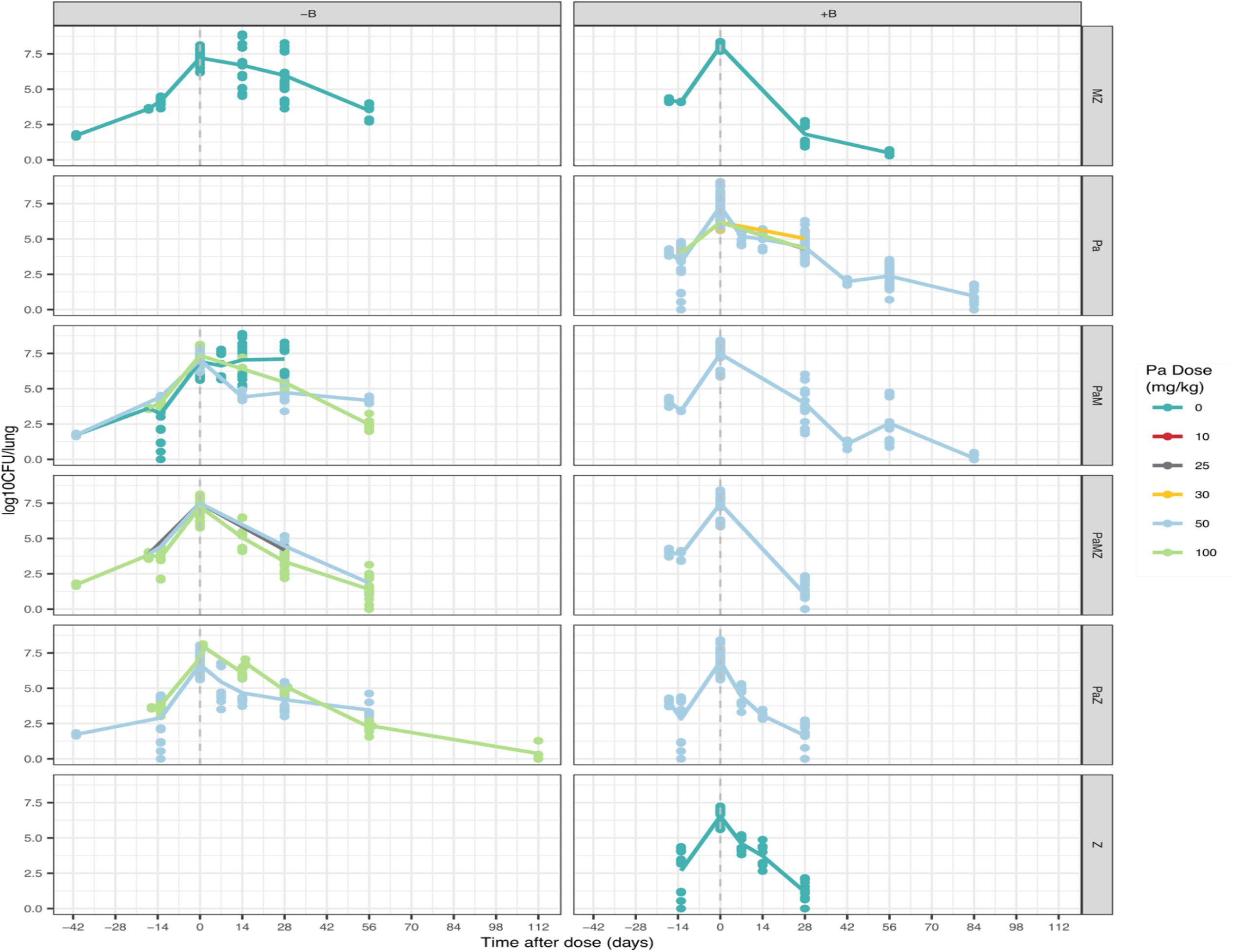
A rich database of high aerosol-infected mice inoculated 14 days prior to treatment with either PaMZ or BPaMZ. Pa had a dose range from 0 – 100mg/kg, while other drugs were at fixed doses. B 25 mg/kg, M 100 mg/kg and Z 150 mg/kg.

**Figure 3.**
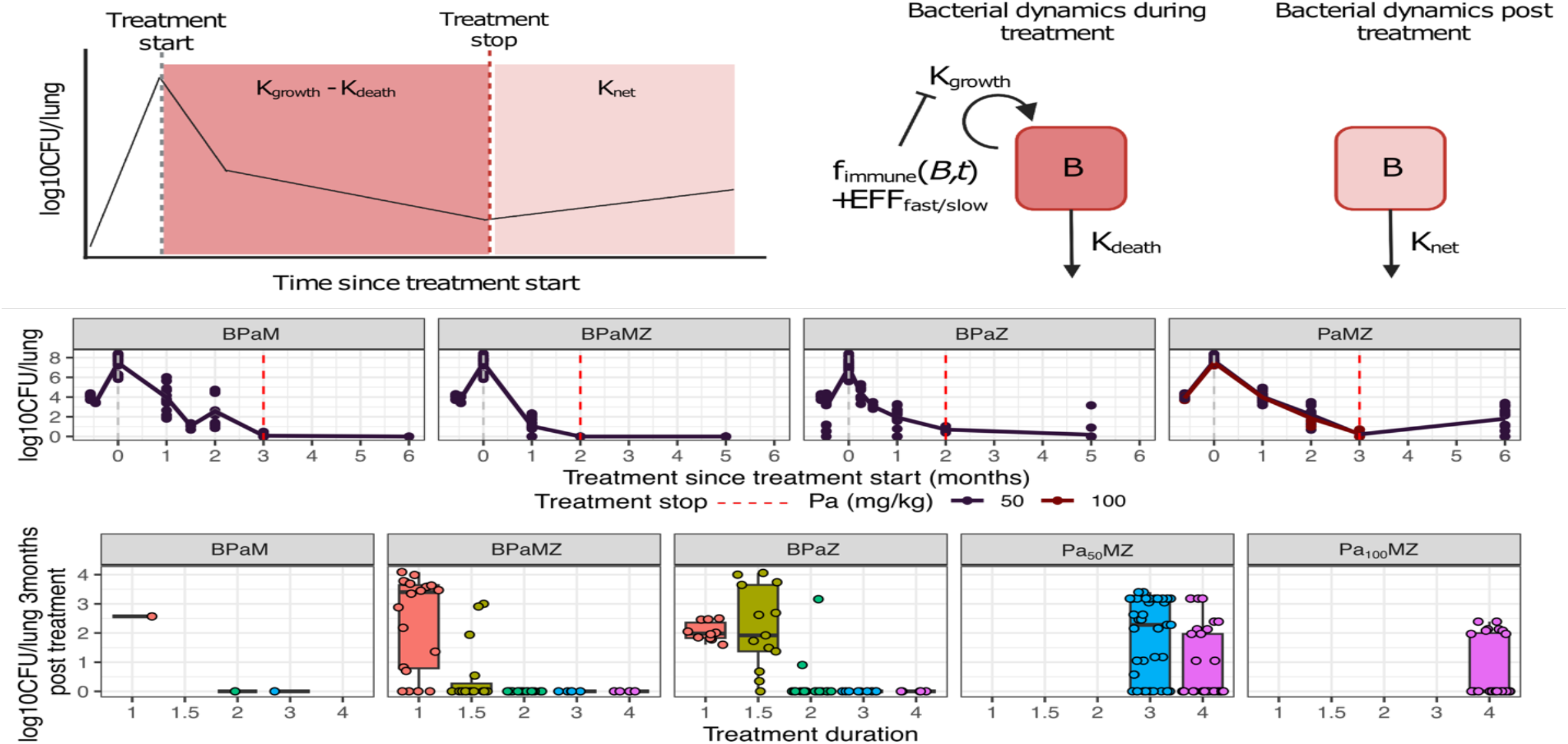
(a) Schematic for modeling mouse CFU during and after treatment. (b) Examples of full mouse CFU profiles for BPaM, BPaMZ, BPaZ and PaMZ during and 3 months post treatment. (c) All available mouse relapse data 3 months after treatment ended. Treatment durations ranged from 1-4 months in mice.

### Estimating fast and slow bacterial growth phases help capture TB persister states

To account for persister bacteria, we assumed the population of fast-replicating bacteria dominated in mice in the first 6 weeks post-inoculation (28 days post-treatment) and the slow-replicating bacteria dominated after^12^. Slow-replicating bacteria were assumed to be less treatment sensitive and thus more reflective of the long-term treatment outcome with Phase IIb/III trials. Our model PK-PD captured this well, with EC50 values of Pa and M showing large decreases in potency from fast to slow growth phase (7.67-fold and 201.7-fold respectively) and B and Z having a small decrease in potency (1.1-fold and 1.27-fold respectively) (Table 2, Figure S2a). This is in line with our current understanding of B and Z having good sterilizing potential in more persister TB states^13,14^.

**Table 2.**
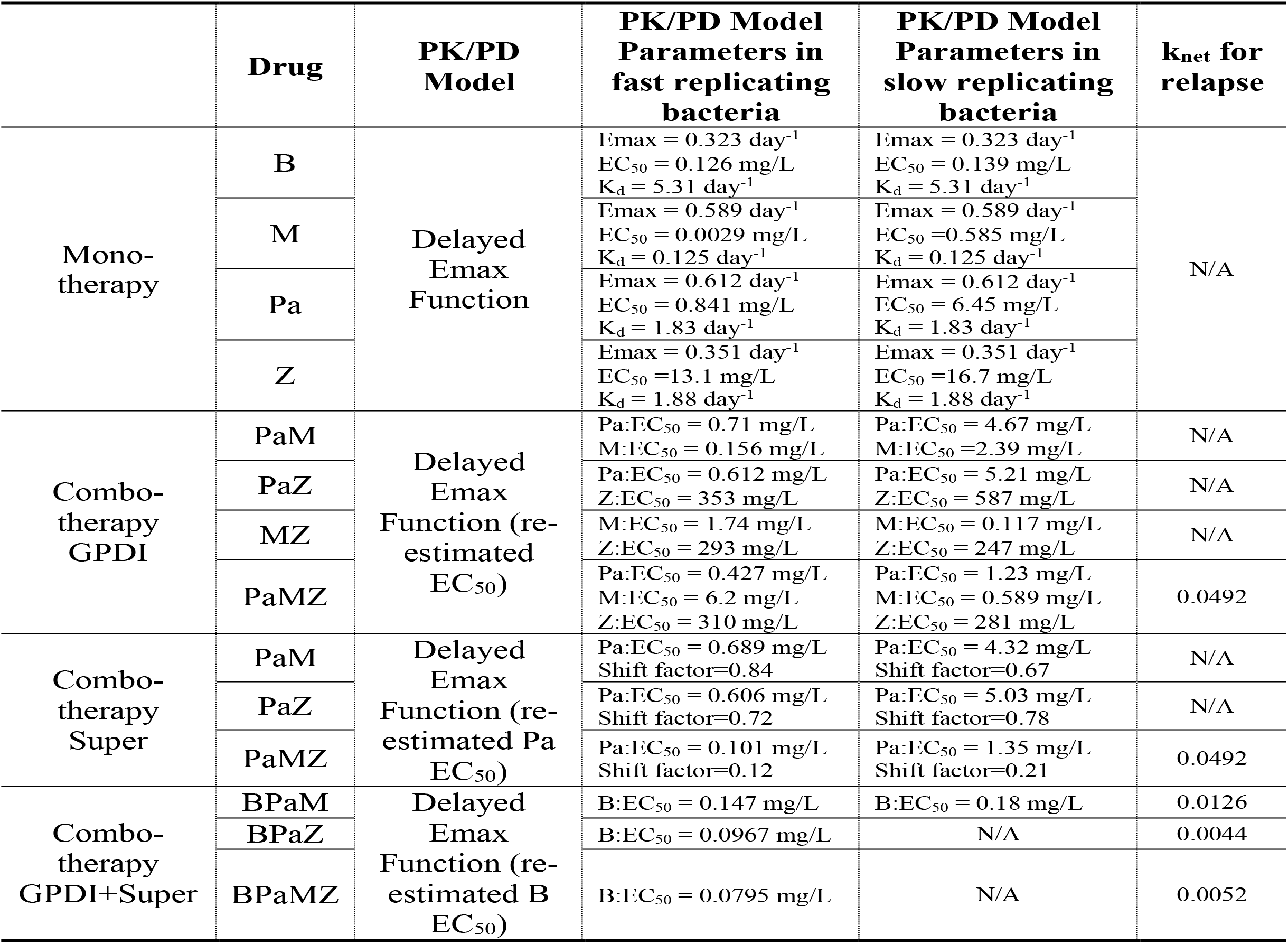
Mouse PK/PD models parameter in fast replicating, slow replicating bacteria and relapse phase using monotherapy and combinations.

| | Drug | PK/PD Model | PK/PD Model Parameters in fast replicating bacteria | PK/PD Model Parameters in slow replicating bacteria | $k_{net}$ for relapse |
| --- | --- | --- | --- | --- | --- |
| Mono-therapy | B | Delayed Emax Function | $E_{max} = 0.323 \text{ day}^{-1}$<br>$EC_{50} = 0.126 \text{ mg/L}$<br>$K_d = 5.31 \text{ day}^{-1}$ | $E_{max} = 0.323 \text{ day}^{-1}$<br>$EC_{50} = 0.139 \text{ mg/L}$<br>$K_d = 5.31 \text{ day}^{-1}$ | N/A |
| | M | | $E_{max} = 0.589 \text{ day}^{-1}$<br>$EC_{50} = 0.0029 \text{ mg/L}$<br>$K_d = 0.125 \text{ day}^{-1}$ | $E_{max} = 0.589 \text{ day}^{-1}$<br>$EC_{50} = 0.585 \text{ mg/L}$<br>$K_d = 0.125 \text{ day}^{-1}$ | |
| | Pa | | $E_{max} = 0.612 \text{ day}^{-1}$<br>$EC_{50} = 0.841 \text{ mg/L}$<br>$K_d = 1.83 \text{ day}^{-1}$ | $E_{max} = 0.612 \text{ day}^{-1}$<br>$EC_{50} = 6.45 \text{ mg/L}$<br>$K_d = 1.83 \text{ day}^{-1}$ | |
| | Z | | $E_{max} = 0.351 \text{ day}^{-1}$<br>$EC_{50} = 13.1 \text{ mg/L}$<br>$K_d = 1.88 \text{ day}^{-1}$ | $E_{max} = 0.351 \text{ day}^{-1}$<br>$EC_{50} = 16.7 \text{ mg/L}$<br>$K_d = 1.88 \text{ day}^{-1}$ | |
| Combo-therapy GPDI | PaM | Delayed Emax Function (re-estimated $EC_{50}$) | Pa:$EC_{50} = 0.71 \text{ mg/L}$<br>M:$EC_{50} = 0.156 \text{ mg/L}$ | Pa:$EC_{50} = 4.67 \text{ mg/L}$<br>M:$EC_{50} = 2.39 \text{ mg/L}$ | N/A |
| | PaZ | | Pa:$EC_{50} = 0.612 \text{ mg/L}$<br>Z:$EC_{50} = 353 \text{ mg/L}$ | Pa:$EC_{50} = 5.21 \text{ mg/L}$<br>Z:$EC_{50} = 587 \text{ mg/L}$ | N/A |
| | MZ | | M:$EC_{50} = 1.74 \text{ mg/L}$<br>Z:$EC_{50} = 293 \text{ mg/L}$ | M:$EC_{50} = 0.117 \text{ mg/L}$<br>Z:$EC_{50} = 247 \text{ mg/L}$ | N/A |
| | PaMZ | | Pa:$EC_{50} = 0.427 \text{ mg/L}$<br>M:$EC_{50} = 6.2 \text{ mg/L}$<br>Z:$EC_{50} = 310 \text{ mg/L}$ | Pa:$EC_{50} = 1.23 \text{ mg/L}$<br>M:$EC_{50} = 0.589 \text{ mg/L}$<br>Z:$EC_{50} = 281 \text{ mg/L}$ | 0.0492 |
| Combo-therapy Super | PaM | Delayed Emax Function (re-estimated Pa $EC_{50}$) | Pa:$EC_{50} = 0.689 \text{ mg/L}$<br>Shift factor=0.84 | Pa:$EC_{50} = 4.32 \text{ mg/L}$<br>Shift factor=0.67 | N/A |
| | PaZ | | Pa:$EC_{50} = 0.606 \text{ mg/L}$<br>Shift factor=0.72 | Pa:$EC_{50} = 5.03 \text{ mg/L}$<br>Shift factor=0.78 | N/A |
| | PaMZ | | Pa:$EC_{50} = 0.101 \text{ mg/L}$<br>Shift factor=0.12 | Pa:$EC_{50} = 1.35 \text{ mg/L}$<br>Shift factor=0.21 | 0.0492 |
| Combo-therapy GPDI+Super | BPaM | Delayed Emax Function (re-estimated B $EC_{50}$) | B:$EC_{50} = 0.147 \text{ mg/L}$ | B:$EC_{50} = 0.18 \text{ mg/L}$ | 0.0126 |
| | BPaZ | | B:$EC_{50} = 0.0967 \text{ mg/L}$ | N/A | 0.0044 |
| | BPaMZ | | B:$EC_{50} = 0.0795 \text{ mg/L}$ | N/A | 0.0052 |

### Pharmacodynamic interactions reveal antagonism but overall additive effects in combinations

Depending on the amount of data available, either the general pharmacodynamics drug interaction model (GPDI) or an empirical approach (SUPER) were used to quantify pharmacodynamic drug-drug interactions for pairwise and 3-way combinations with Pa, M and Z (Figure S1). GPDI was used only when sufficient dose ranges were available both in monotherapy pairwise and 3-way combinations to predict the effect of each drug. The SUPER approach on the other hand, only required the determination of the exposure-response relationship of the “super” drug whereas the existing drug combination could remain at fixed doses. GPDI models which estimated the contribution of each drug to the overall combined effect of a drug combination showed PaM and PaZ combinations showed adding either M or Z to a Pa improved Pa potency in both fast and slow growth phases. This was similarly observed when we applied SUPER to understand the effect of adding either M or Z to Pa, with Pa EC50 decreasing in combinations compared to monotherapy (Table 2). Adding both M and Z to Pa (PaMZ) gave the most significant increase in Pa potency (1.96-fold decrease in EC50 using GPDI, shift factor 0.12 using SUPER). This suggests adding MZ to Pa provides both a synergistic as well as additive effect to Pa. Interestingly, while M as monotherapy had a large drop in potency from fast to slow growth phase, M as a combination in either PaM or PaMZ was found to be more potent in slow-replicating bacteria than fast-replicating bacteria. This suggests a possible pharmacologic effect that M needs to be combined with another drug to exert its sterilizing activity. Interestingly, B did not show much PD drug interactions and did not change much in potency between fast and slow replicating states (Table 2).

### A combination GPDI and SUPER model to estimate bedaquiline’s effect

B was only tested at a fixed dose of 25 mg/kg in mice. To tease out the individual contribution of B to the PaMZ regimen, we combined the GPDI and SUPER method, using B as the super drug, and either PaZ or PaMZ as the backbone regimen (Table 2). This method assumes B does not have a DDI with the backbone regimen and estimates the individual effect of B being added to the regimen. The potency of B in combination BPaZ and BPaMZ decreased in comparison to monotherapy, indicating synergism between B and both PaZ and PaMZ.

### Visual predictive checks (VPCs) indicate model reliability

All models using either GPDI or SUPER were evaluated using visual predictive checks (VPCs) of 200 simulations which showed the observed data was consistently within the 95% prediction interval of the bacterial numbers in the final PK/PD models for all three approaches (Figure 3).

### Validation and prediction of long-term phase IIb & III study outcomes

We first validated our translational predictions with short term early bactericidal activity (EBA) studies. Using PK-PD indices from the fast replicating growth phase in our mouse studies, we predicted EBA in clinical studies NC-001 and NC-003 well as observed with the majority of the clinical data points lying within the model’s 95% prediction interval ^15,16^ (Supplemental Table 3). With increased confidence from short-term treatment, we used PK-PD relationships identified in both fast- and slow-replicating bacteria to predict long-term treatment (Figure 3). The long-term treatment outcome of interest was the proportion of patients with undetectable CFU post-treatment. This proportion was predicted by simulating bacterial regrowth post-treatment to determine the proportion of patients that would have undetectable CFU, defined by CFU < 1, using different treatment durations. Each study was simulated according to its own baseline demographic information. The proportion of patients with undetectable CFU was predicted well in BPaZ and PaMZ regimens (Figure 4a) with the actual clinical trial proportions lying close to the median of the model predictions. BPaMZ from clinical observations (NC-005) were also captured within the lower bound of the model prediction interval. This over-prediction is possible because patients in NC-005 were MDR-TB patients rather than drug-sensitive TB patients as with the other trials.

**Table 3.**
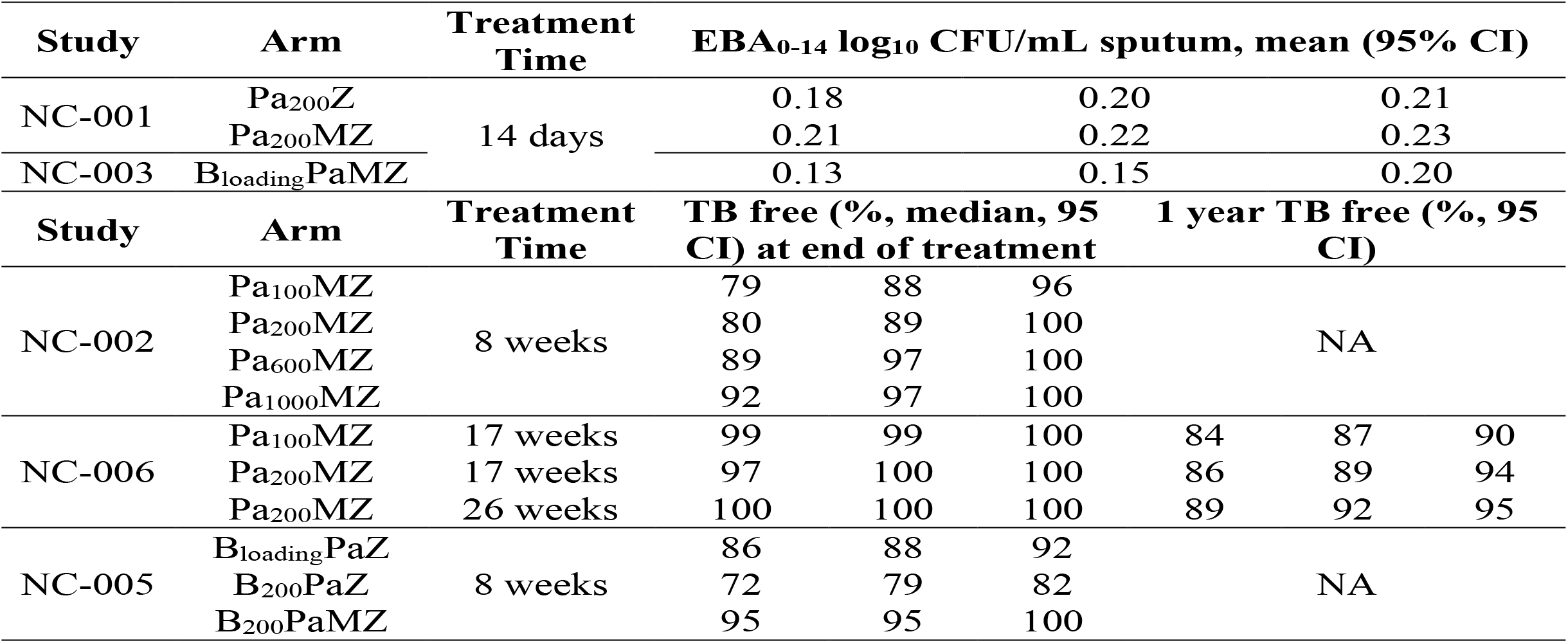
Human outcome predictions for short- and long-term treatment of PaMZ and BPaMZ regimens.

**Figure 4.**
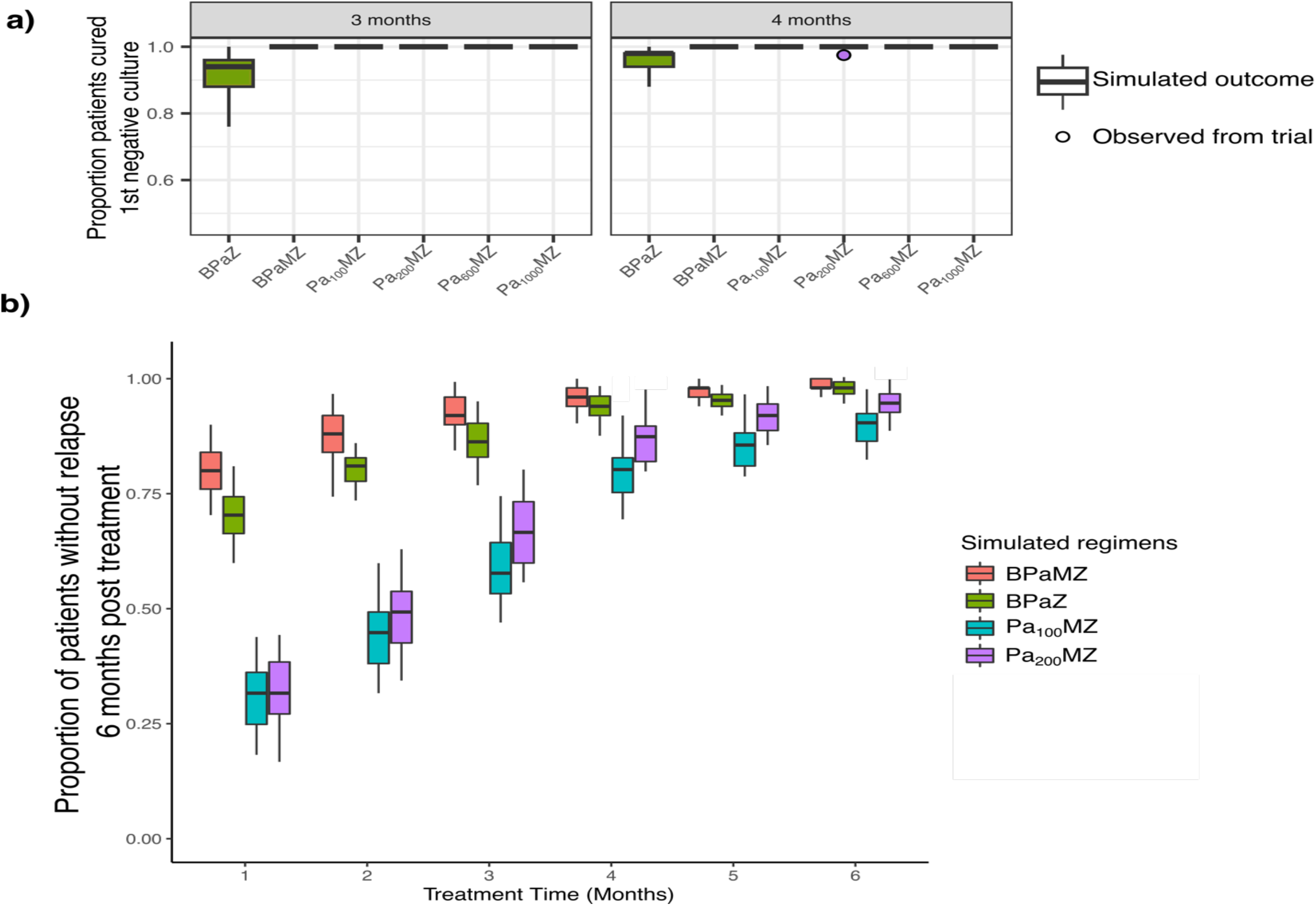
(a) Ranking regimens by cure rate at 3- and 4-month durations. Predicted human outcome (proportion of CFU negative patients, box plots) overlapped with clinical observations (points). (b) Regimens rank-ordering based on the proportion of non-relapse patients 6 months after different treatment durations.

### Drug combination rank-ordering based on treatment effect

With confidence in our model, we then decided to rank the regimens based on how quickly they were predicted to reach a proportion of patients with undetectable CFU of 1 in the span of 6 months. Undetectable CFU suggests a low probability of relapse as all bacteria should have been eliminated. Baseline demographic information in NC-002, NC-005, and NC-006 were pooled and used to simulate all 8 regimens for a fair comparison. Pa_1000_MZ had the fastest response at 1.5 months, followed by BPaMZ at 2 months. All regimens showed 100% of patients with undetectable CFU, at 4 months (Figure 4a). Our model thus points toward either a high dose pretomanid > 600mg or the current recommended dose with added B as a 4^th^ drug for a shorter treatment duration of 4 months.

### Predicting relapse 6 months post-treatment showed B containing regimens were superior

To further differentiate a superior regimen, we extended our platform further to predict relapse 6 months post-treatment. We simulated a range of treatment durations from 1-6 months using regimens in NC-006 (Pa_100_MZ and Pa_200_MZ) and SimpliciTB (BPaMZ) to simulate the proportion of non-relapse patients 6 months after treatment ended. B-containing regimens were superior to non-B containing regimens with a higher proportion of patients with no relapse with 1-4 months of treatment (Figure 4b). It is worth noting, however, that our model had a small underprediction of the proportion of relapse patients as seen with actual relapse data for Pa_100_MZ and Pa_200_MZ. The simulation of clinical log10CFU during and post treatment can be found in Figure S5. Based on these results, we predict that adding B will improve the PaMZ regimen, indicating the SimpliciTB trial is likely to be successful if safety events are not limiting.

## Discussion

We extended our translational platform from predicting short-term monotherapy clinical outcomes to include long-term combination treatment outcomes bacteria burden and proportion of relapse. Our platform was able to predict short-term outcomes as EBA as well as long-term outcomes proportion of patients with undetectable CFU and relapse 6 months post-treatment.

Our model was able to predict the proportion of patients with undetectable CFU well. While there was no significant difference in cure rate at 4 months of treatment between high dose Pa regimens 600mg or lager and regimens with B added, adding B as a 4^th^ drug is likely to be preferable due to the lower rate of relapse 6 months post-treatment.

When predicting relapse 6 months post-treatment, despite a small amount of underprediction, we were still able to differentiate efficacy between treatment regimens making our platform efficacious in ranking and prioritizing regimens for testing.

Although the murine TB model lacks lung lesions, it is still an important preclinical model due to the ease of handling and relatively low procurement and housing costs. Our work has shown the importance of modeling both a fast and slow bacterial growth phase as each growth phase reflects a translatable drug in vivo potency from mice to human in short and long term outcomes respectively ^17–19^.

Our bacterial dynamics model was also important in helping to predict initial bacterial growth and death after inoculation and during treatment. However, as bacterial regrowth data after 3 months of treatment was sparse, we were not able to train a full bacterial dynamics model with confidence for regrowth. We thus used K_net_ as a composite measure of both bacterial growth and death instead to describe the net bacteria regrowth. In comparison to non-B regimens, K_net_ was much slower with B onboard (Table 2). This could be a result of the sterilizing activity ^20^ and long half-life ^21^ of B. Moreover, in B-containing regimens, with Z onboard, K_net_ was even smaller, indicating B in combination with Z had even better sterilizing activity.

Another key factor to our successful translation of human clinical outcomes was being able to estimate the individual contribution of each drug in combination or the effect of a backbone regimen on a newly added drug. This allowed us to estimate the individual contribution of adding B and increasing the dose of Pa in different drug regimens, allowing a quantifiable way to predict outcomes for new regimens.

We applied GPDI and SUPER approaches in the current study based on data availability (Fig S2a). GPDI was used only when sufficient dose ranges were available both in monotherapy pairwise and 3-way combinations to predict the effect of each drug. The SUPER approach on the other hand, only required the determination of the exposure-response relationship of the dose-ranging drug whereas the backbone drug combination could remain at fixed doses. In our exploration with PaMZ, we found no significant difference in model predictions using both GPDI and SUPER to fit mouse CFU data (Supplementary Fig 2b). To better account for the individual effect of B, we thus modified the GPDI method to exploit the developed GPDI model in PaMZ combinations, allowing us to capture the BPaMZ mouse CFU profile well (Figure 3). Even though for each drug in PaMZ, INT was greater than 1 meaning antagonism, the combined effect of PaMZ was still superior to Pa monotherapy. Although synergism is preferable in drug combinations, many antibiotic combinations for TB treatment tend to be antagonistic in nature but have an overall additive effect, resulting in desirable clinical outcomes overall. Therefore, using proposed translational platform could help select more promising drug combinations even though the components could be antagonistic.

In novel drug combination development pipeline, new drugs are usually introduced as an additional drug combined with an existing backbone regimen^22^. To design efficient animal experiments, we recommend testing the drug of interest with at least three different doses in each combination.

Our proposed translational platform successfully translated findings in preclinical murine models to predict long-term human clinical outcomes. This serves as a useful tool in rank ordering regimens based on both cure and relapse rates, helping us to prioritize the regimens most likely to shorten treatment duration while demonstrating non-inferiority to the current standard of care.

## Methods

### Mouse PK-PD model development

Mouse PK-PD models were developed on top of a previously published bacterial infection model that describes bacteria growth, death, and adaptive immune effect without drug treatment ^12^. Effect of drug combinations were estimated using the general pharmacodynamics drug interaction model (GPDI) approach and an empirical approach (SUPER), based off the availability and abundance of data in murine model. Drug effect of combinations were evaluated in both fast- and slow-replicating bacteria. Additive, proportional and combined error models were tested to describe residual error for the mouse PK/PD models.

### Mice relapse model

A simplified baseline model measuring only the net growth rate constant (K_net_) of bacteria was used to describe mice relapse. Net growth of bacteria after treatment stopped was calculated. Treatment duration and drug combination were also added as covariates to account for differences in experiments.

### Translational model development for short-term EBA prediction

The established mouse PK/PD relationships in fast-replicating bacteria only in combinations PaZ, PaMZ, and BPaMZ was translated to human by protein binding correction followed by prediction of clinical outcomes of EBA studies NC-001 and NC-003. Clinical outcomes included average daily drop of log_10_ CFU values and longitudinal log_10_ CFU change over 14-day’s treatment. Clinical outcomes were simulated by predicting the CFU counts in TB patients using the translated PK-PD relationship determined in the murine model. Two hundred simulations were conducted for each drug.

### Translational model for long-term phase 2b and phase 3 prediction and simulation

Drug efficacy in fast-replicating bacteria was used to predict clinical early bactericidal activity up to 14 days (Figure S4). Then PK-PD relationships were shifted to that determined in slow-replicating bacteria to predict CFU counts in patients in post 14-day’s treatment. Existing phase 2b and 3 trial regimens (NC-002, NC-005, NC-006) were included in the simulation first. Human PK of Pa, M Z and B were simulated using patients’ demographic data at baseline and detailed in supplemental material. CFU counts change over time were simulated using patient’s individual CFU counts at baseline. Five hundred simulations were conducted for each regimen. Based on simulated CFU values, proportions of negative CFU (bacteria number less than 1) and proportion of relapse 6-month after treatment stops were included as clinical outcome. In addition to existing trials, we add several regimens that has not been tested in human (Figure 3 and Figure 4) with multiple dosing regimens of Pa or B at different durations from 1 to 6 months. Patients were pulled population from existing NC-002, NC-005, NC-006, assuming that patient’s demographic information were normally distributed. Five hundred simulations were conducted. In addition, in each regimen (both in existing trial or non-tested in human), after treatment stops, we continue to simulate an additional 6 months and calculate the proportion of patients that have bacterial number larger than 1 as “relapse”. Proportion of relapse patients were calculated and compared between regimens.

---

**Figure S1.**
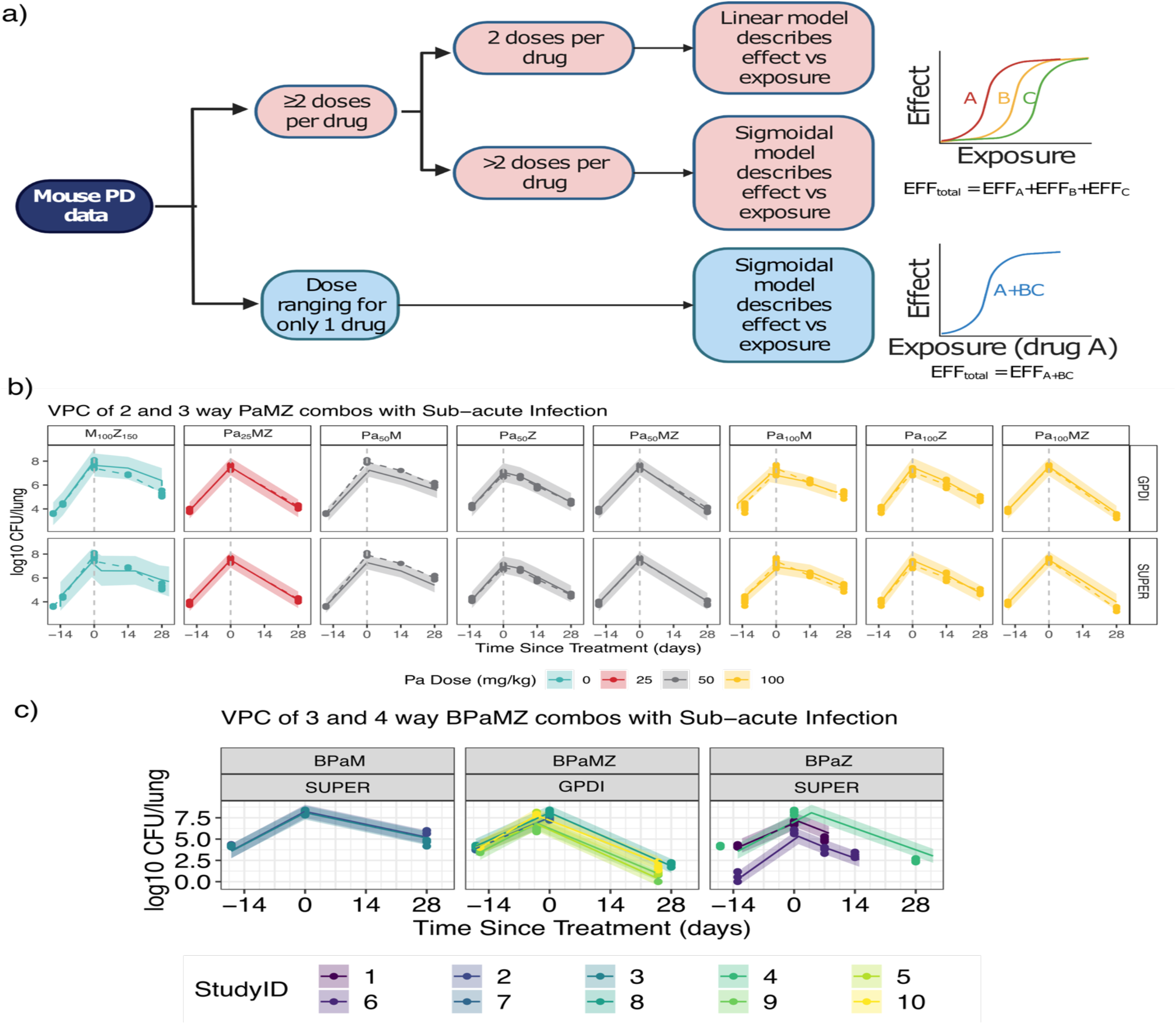
(a) Illustration of GPDI and SUPER approach based on data availability. (b) visual predictive checks of both GPDI and SUPER methods show models describe PaMZ data well. (c) BPaMZ was similarly described well by both SUPER and GPDI methods.

**Figure S2.**
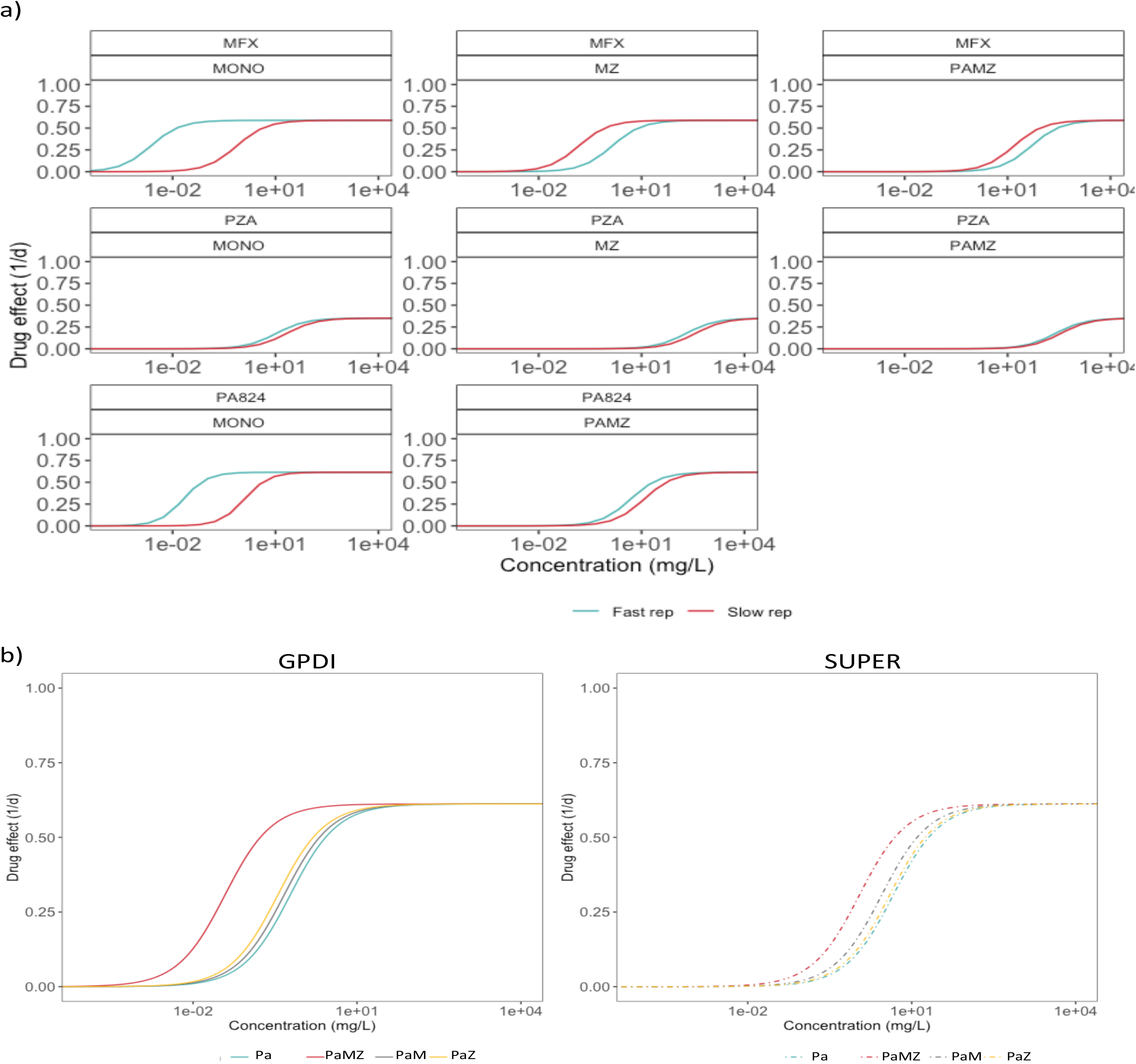
Drug potency of Pa, M and Z in different PaMZ combinations using GPDI approach (a) in both fast- and slow-replicating bacteria in mice. (b) drug potency of Pa in different PaMZ combinations using either GPDI or SUPER approach

**Figure S3.**
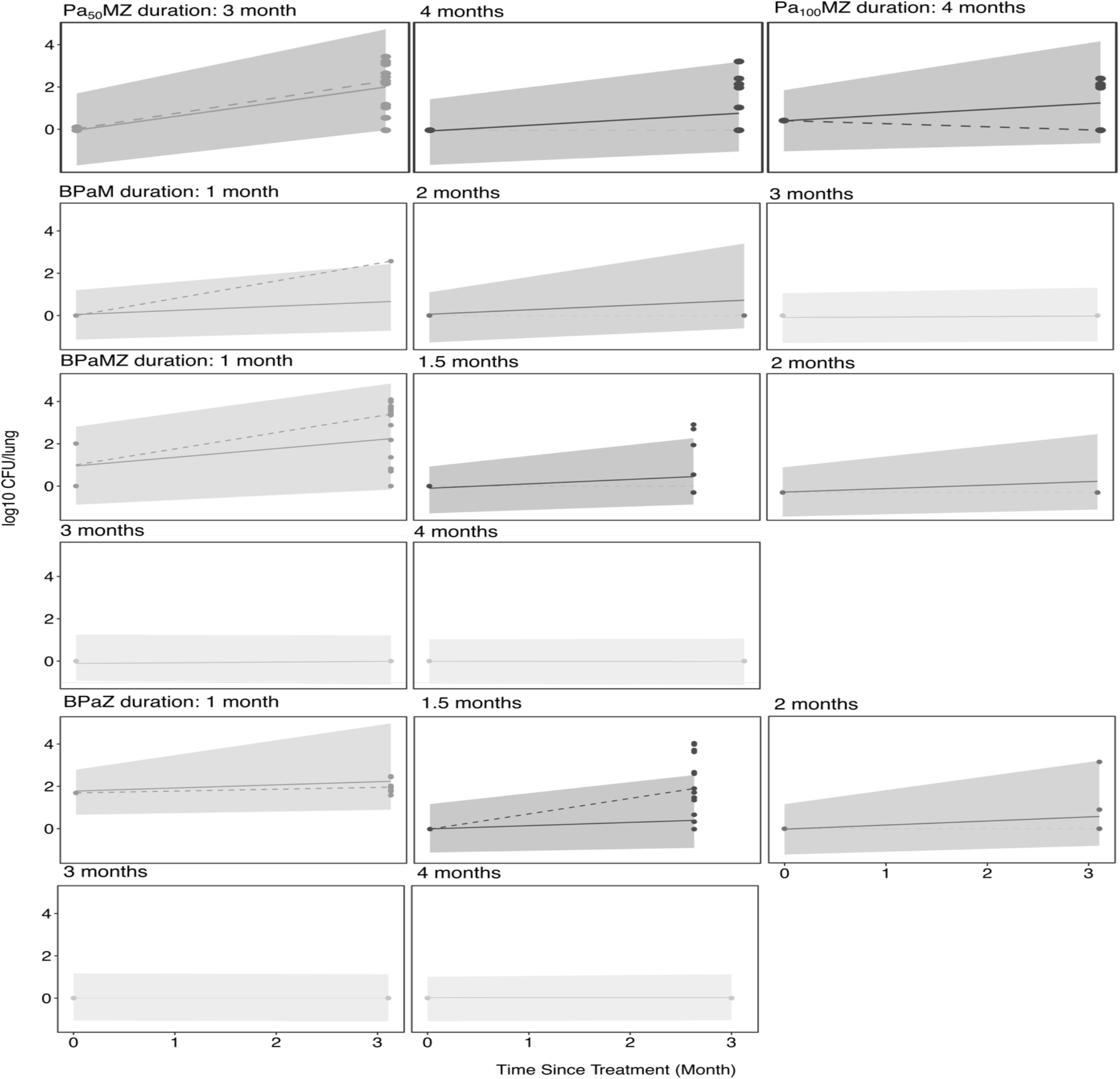
Mice relapse modeling validation after treatment of different PaMZ and BPaMZ combinations after various treatment durations.

**Figure S4.**
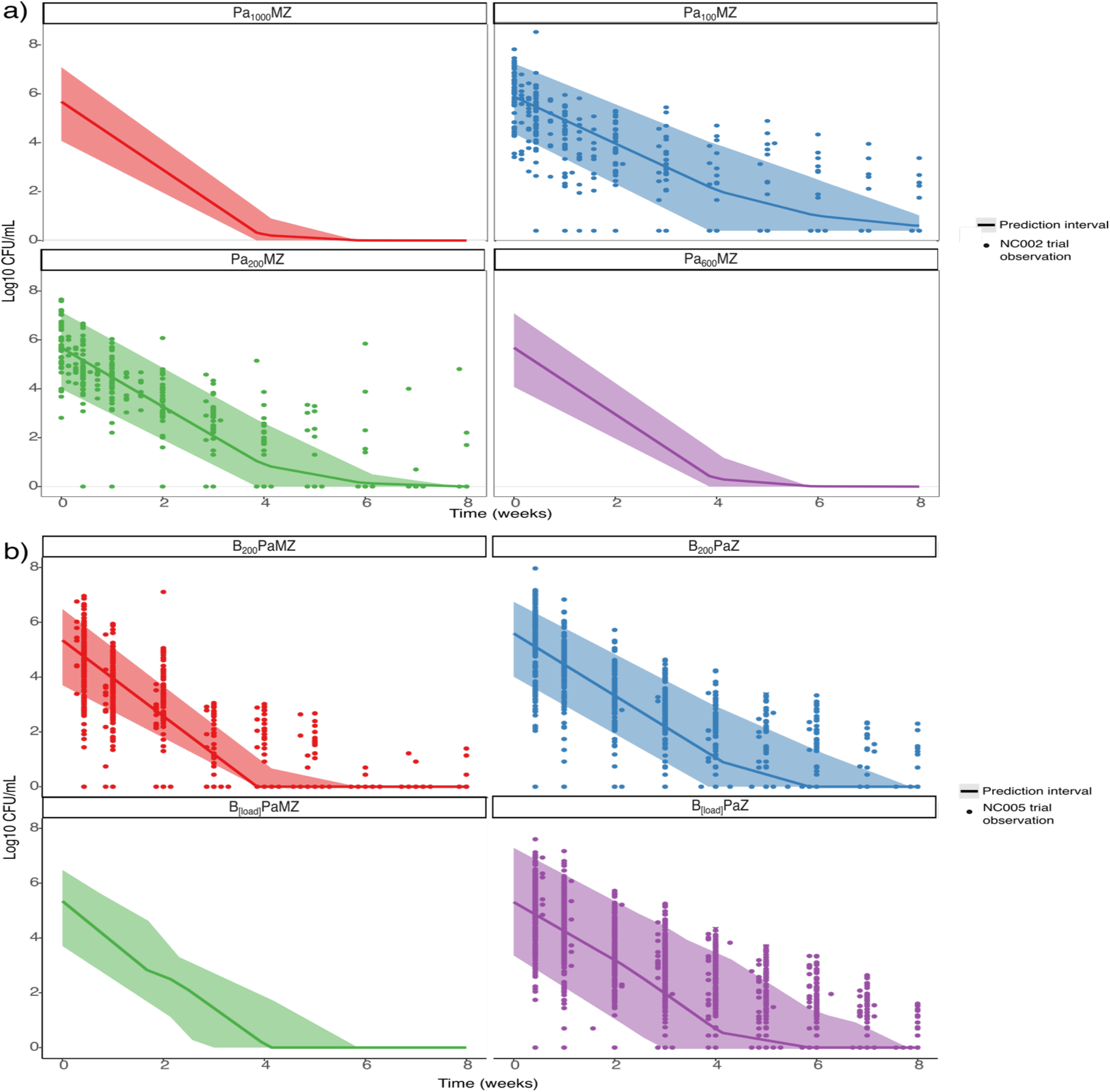
Early bactericidal activity predictions of PaMZ regimens in NC-002 trials (a), and BPaMZ regimens in NC-005 trials (b).

**Figure S5.**
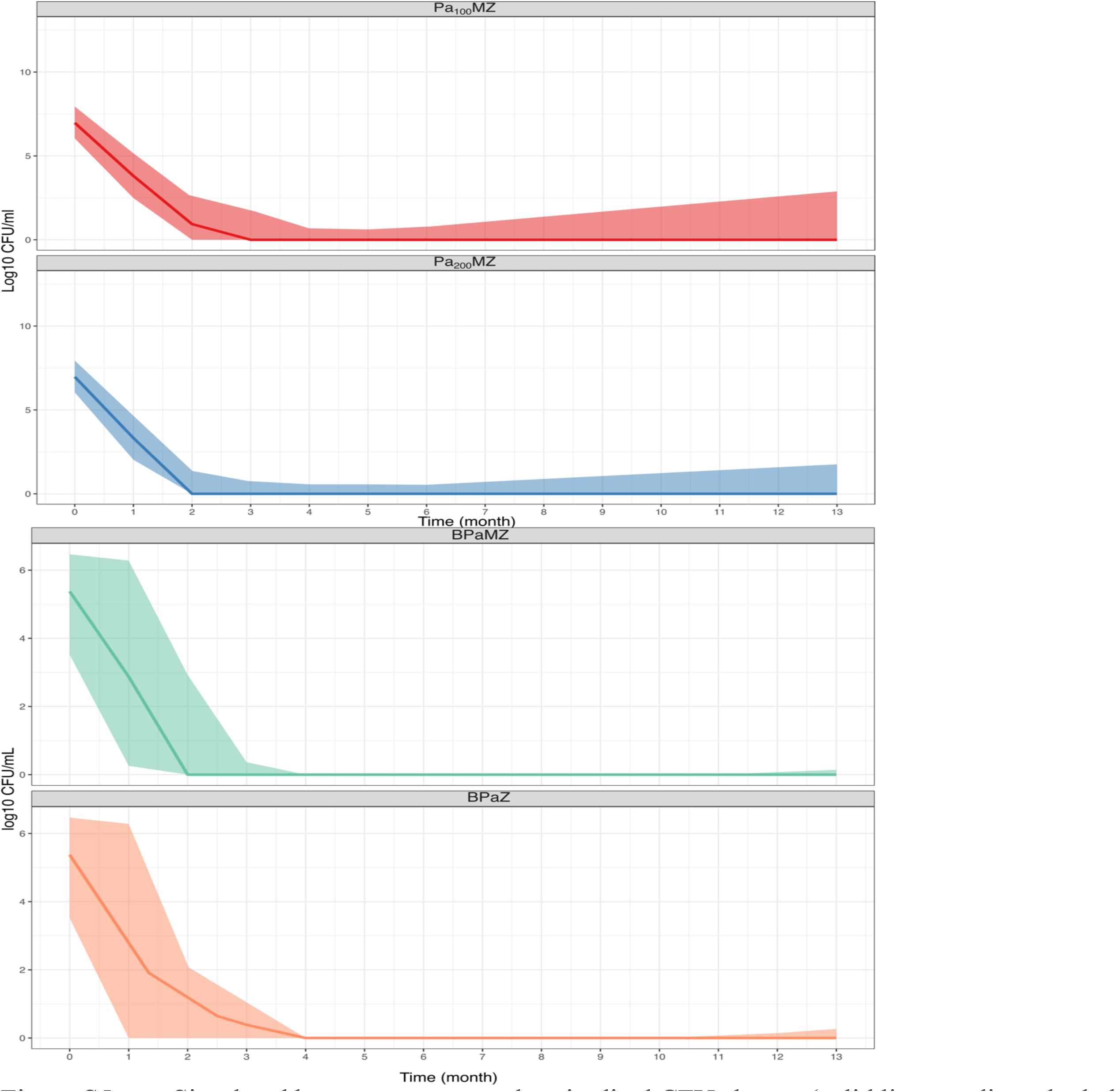
Simulated human outcome as longitudinal CFU change (solid line: median; shaded area: 95%CI of simulation) during treatment and bacteria re-growth in treatment of PaMZ (top) and BPaMZ (bottom). Treatment duration for all regimens were fixed at 4-month.

**Figure S6.**
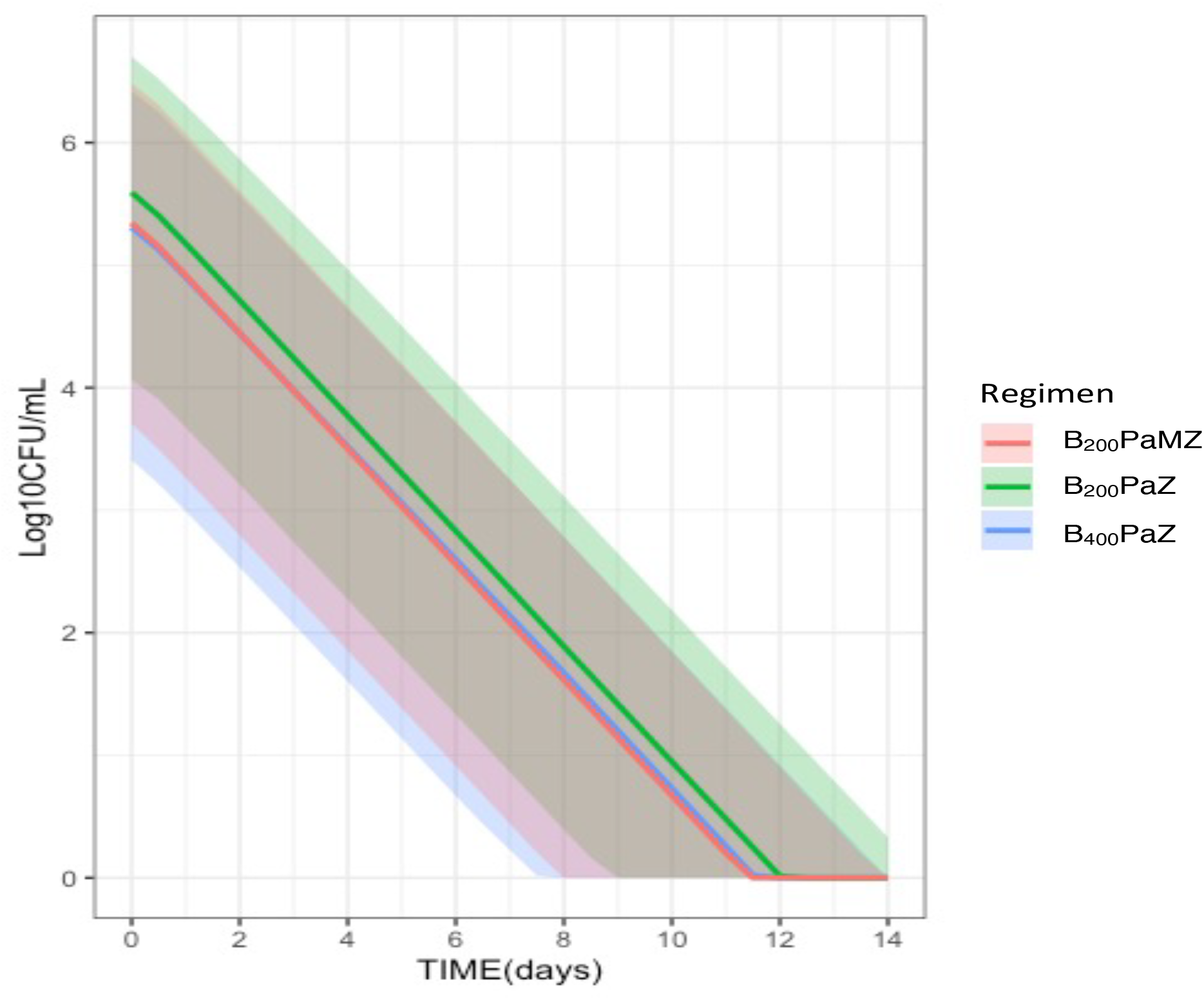
Simulation of BPaZ in human at various dosing regimens for B by PK/PD relationship determined in mice using only SUPER approach.

**Table S1.**
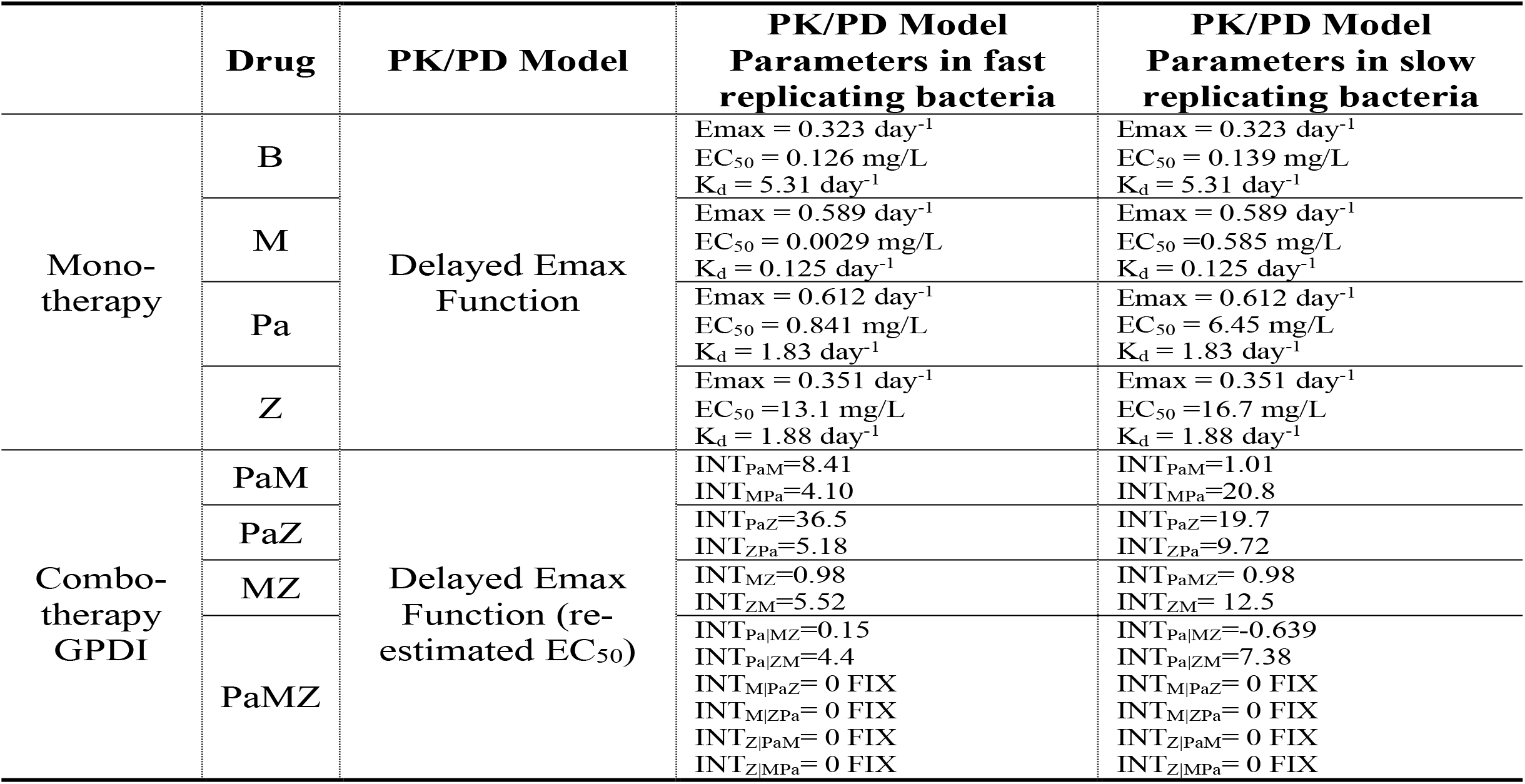
Drug-drug interaction in PaMZ combinations analyzed by GPDI approach in mice.

|  | Drug | PK/PD Model | PK/PD Model<br>Parameters in fast<br>replicating bacteria | PK/PD Model<br>Parameters in slow<br>replicating bacteria |
| --- | --- | --- | --- | --- |
| Mono-<br>therapy | B | Delayed Emax<br>Function | $E_{max} = 0.323 \text{ day}^{-1}$<br>$EC_{50} = 0.126 \text{ mg/L}$<br>$K_d = 5.31 \text{ day}^{-1}$ | $E_{max} = 0.323 \text{ day}^{-1}$<br>$EC_{50} = 0.139 \text{ mg/L}$<br>$K_d = 5.31 \text{ day}^{-1}$ |
| | M | | $E_{max} = 0.589 \text{ day}^{-1}$<br>$EC_{50} = 0.0029 \text{ mg/L}$<br>$K_d = 0.125 \text{ day}^{-1}$ | $E_{max} = 0.589 \text{ day}^{-1}$<br>$EC_{50} = 0.585 \text{ mg/L}$<br>$K_d = 0.125 \text{ day}^{-1}$ |
| | Pa | | $E_{max} = 0.612 \text{ day}^{-1}$<br>$EC_{50} = 0.841 \text{ mg/L}$<br>$K_d = 1.83 \text{ day}^{-1}$ | $E_{max} = 0.612 \text{ day}^{-1}$<br>$EC_{50} = 6.45 \text{ mg/L}$<br>$K_d = 1.83 \text{ day}^{-1}$ |
| | Z | | $E_{max} = 0.351 \text{ day}^{-1}$<br>$EC_{50} = 13.1 \text{ mg/L}$<br>$K_d = 1.88 \text{ day}^{-1}$ | $E_{max} = 0.351 \text{ day}^{-1}$<br>$EC_{50} = 16.7 \text{ mg/L}$<br>$K_d = 1.88 \text{ day}^{-1}$ |
| Combo-therapy<br>GPDI | PaM | Delayed Emax<br>Function (re-estimated $EC_{50}$) | $INT_{PaM}=8.41$<br>$INT_{MPa}=4.10$ | $INT_{PaM}=1.01$<br>$INT_{MPa}=20.8$ |
| | PaZ | | $INT_{PaZ}=36.5$<br>$INT_{ZPa}=5.18$ | $INT_{PaZ}=19.7$<br>$INT_{ZPa}=9.72$ |
| | MZ | | $INT_{MZ}=0.98$<br>$INT_{ZM}=5.52$ | $INT_{PaMZ}=0.98$<br>$INT_{ZM}=12.5$ |
| | PaMZ | | $INT_{Pa\vert MZ}=0.15$<br>$INT_{Pa\vert ZM}=4.4$ | $INT_{Pa\vert MZ}=-0.639$<br>$INT_{Pa\vert ZM}=7.38$ |
| | | | $INT_{M\vert PaZ}=0 \text{ FIX}$<br>$INT_{M\vert ZPa}=0 \text{ FIX}$<br>$INT_{Z\vert PaM}=0 \text{ FIX}$<br>$INT_{Z\vert MPa}=0 \text{ FIX}$ | $INT_{M\vert PaZ}=0 \text{ FIX}$<br>$INT_{M\vert ZPa}=0 \text{ FIX}$<br>$INT_{Z\vert PaM}=0 \text{ FIX}$<br>$INT_{Z\vert MPa}=0 \text{ FIX}$ |

